# Metabolic and functional connectivity provide unique and complementary insights into cognition-connectome relationships

**DOI:** 10.1101/2021.09.06.459214

**Authors:** Katharina Voigt, Emma X. Liang, Bratislav Misic, Phillip G.D. Ward, Gary F. Egan, Sharna D. Jamadar

**Author notes:** Corresponding author: Dr Katharina Voigt, Turner Institute for Brain and Mental Health, Monash Biomedical Imaging, 770 Blackburn Road, Clayton, VIC 3800, Australia.

## Abstract

A major challenge in current cognitive neuroscience is how functional brain connectivity gives rise to human cognition. Functional magnetic resonance imaging (fMRI) describes brain connectivity based on cerebral oxygenation dynamics (hemodynamic connectivity), whereas [18 F]-fluorodeoxyglucose functional positron emission tomography (FDG-fPET) describes brain connectivity based on cerebral glucose uptake (metabolic connectivity), each providing a unique characterisation of the human brain. How these two modalities differ in their contribution to cognition and behaviour is unclear. We used simultaneous resting-state FDG-fPET/fMRI to investigate how hemodynamic connectivity and metabolic connectivity relate to cognitive function by applying partial least squares analyses. Results revealed that while for both modalities the frontoparietal anatomical subdivisions related the strongest to cognition, using hemodynamic measures this network expressed executive functioning, episodic memory, and depression, while for metabolic measures this network exclusively expressed executive functioning. These findings demonstrate the unique advantages that simultaneous FDG-PET/fMRI has to provide a comprehensive understanding of the neural mechanisms that underpin cognition and highlights the importance of multimodality imaging in cognitive neuroscience research.

## Introduction

The human connectome is a comprehensive map of neural connections, that describes the brain as a complex network of interconnected brain regions (Sporns, 2013; Sporns et al., 2005). Non-invasive neuroimaging methods provide us with the opportunity to characterise the functionality of brain connectivity on multiple levels (Raichle, 2009). As such, brain connectivity is a multidimensional concept that is defined by its measurement tool. Blood oxygenation level dependent (BOLD) functional magnetic resonance imaging (fMRI) has been the dominant tool to characterise functional brain connectivity, based on the temporal coherence of spontaneous, low-frequency large-amplitude changes in blood oxygenation whilst an individual is at rest (Biswal et al., 1995; Raichle, 2011). BOLD-fMRI provides a haemodynamic-based surrogate measure of neuronal activity at a high spatial and temporal resolution, but is confounded by non-neuronal components (e.g., heart rate, respiration, blood volume; Liu, 2017; Ward et al., 2020). Positron Emission Tomography (PET) scanning using the glucose analogue F18-fluordoxyglucose (FDG) provides the opportunity to characterise metabolic elements of brain connectivity based on cerebral glucose update (Yakushev et al., 2017). In contrast to BOLD-fMRI, FDG-PET is a quantifiable index of neuronal activity capturing cerebral glucose uptake at the synapses. The integration of the two modalities in a simultaneous MR-PET system (Chen et al., 2018; Judenhofer et al., 2008) offers the unique opportunity to undertake multidimensional neuroimaging studies to examine the interaction between hemodynamic and metabolic aspects of brain connectivity. How these elements of human brain connectivity (i.e., hemodynamic and metabolic) individually and jointly contribute to human cognition and behaviour remains a formidable challenge of contemporary cognitive neuroscience.

A central assumption in cognitive neuroscience is that cognitive processes are emergent properties of neural communication, which is predicted by the coherent and flexible oscillatory activity between neural ensembles (Avena-Koenigsberger et al., 2018; Barack and Krakauer, 2021). Two views of how the brain as a neural network relates to cognition have emerged: the *domain-specific* connectome-cognition view and the *global* connectome-cognition view. According to the specific view, connectivity is domain-specific and multiple networks arise for distinct cognitive domains. According to the global view, the overall wiring of connectome gives rise to global cognitive functioning. A single set of connectivity patterns predict cognitive functioning across different domains, such as attention, memory, executive functioning.

Evidence from fMRI research using multivariate analytic approaches to examine brain-behaviour relationships has revealed support for both views (e.g., Goyal et al., 2020; Smith et al., 2015; Ziegler et al., 2013; Zimmermann et al., 2018). In support of the domain-specific connectome-cognition view, Zimmermann and collegues (2018) found unique orthogonal sets of resting-state hemodynamic connectivity clusters that were associated with specific cognitive domains. Inter- and intra-hemispheric resting-state hemodynamic connectivity in the frontoparietal, occipital, temporal, and cingulate areas was negatively associated with processing speed, executing functioning, and working memory. Intelligence was related to a separate set of resting-state hemodynamic connectivity in cortico-cortical and cortico-subcortical networks, such as the caudate and putamen. In contrast, in support of the global view, Smith and others (2015) revealed a single mode of large-scale resting-state hemodynamic connectivity patterns capturing a wide set of behavioural (e.g., intelligence, verbal ability) and demographic variables (e.g., age, sex, income, drug use). This result has recently been replicated by Goyal and others (2020). These studies reveal initial insights into how coherent, low-frequency BOLD fMRI signalling in spatially distinct brain areas (i.e., hemodynamic connectivity) relates to cognition. However, as the BOLD signal represents a proxy of neural activity that is shaped by non-neuronal contributions to the BOLD signal (Liu, 2017; Ward et al., 2020), this considerably restricts our existing understanding of connectome-cognition systems to haemodynamic correlates.

Recent developments in continuous radiotracer delivery and improved PET signal detection of dual-modality magnetic resonance (MR)-PET scanners, has allowed the study of continuous glucose uptake with substantially improved temporal resolution (e.g., 60 seconds or less; Jamadar et al., 2021; Rischka et al., 2018; Villien et al., 2014). This novel method, termed “functional” FDG-PET (FDG-fPET), provides the opportunity to characterise the metabolic connectome beyond previous covariance measures resulting from static PET (Jamadar et al., 2021) and thus, approaches similar within-subject time-course correlational descriptions as exist for BOLD-fMRI hemodynamic connectivity. Using the fPET approach, we recently found that the metabolic FDG-fPET connectome showed moderate similarity with the BOLD-fMRI hemodynamic connectivity at rest, with the highest similarity between functional and metabolic connectivity obtained primarily with the superior and frontoparietal cortical areas (Jamadar et al., 2021). These initial findings suggest the complementary potential of describing the human connectome via fMRI and FDG-fPET. However, how resting-state metabolic connectivity derived from FDG-fPET relates to cognition and how it differs in their predictive ability from BOLD-fMRI hemodynamic connectivity remains unknown.

The present study aimed to investigate whether (1) a single global or multiple distinct connectivity pattern maps onto cognition, and whether (2) the connectome-cognition relationship is different for hemodynamic and metabolic connectivity derived from a novel FDG-fPET methodology (Jamadar et al., 2020; Jamadar et al., 2021). We acquired FDG-fPET data with high temporal resolution of 16s to measure glucose metabolic connectivity, and simultaneously acquired BOLD-fMRI data with a temporal resolution of 2.45s from 26 participants. Participants completed a neuropsychological cognitive test battery, which resulted in 14 cognitive outcome variables indexing cognition across several domains (verbal memory, attention, executive functioning). We used partial least squares (PLS) to map orthogonal patterns of brain-behaviour relationships (Krishnan et al., 2011; McIntosh and Lobaugh, 2004; McIntosh and Misic, 2013) (Figure 1). As evidence for both connectome-cognition views has been reported for fMRI data (e.g., Goyal et al., 2020; Smith et al., 2015; Ziegler et al., 2013; Zimmermann et al., 2018), we undertook and exploratory analysis to investigate how hemodynamic and metabolic connectomes map onto cognition. However, as hemodynamic and metabolic connectomes have been shown to reveal distinct connectivity patterns (Jamadar et al., 2021), we hypothesised that both would provide a unique, but complementary insight, into the connectome-cognition relationship.

**Figure 1.**
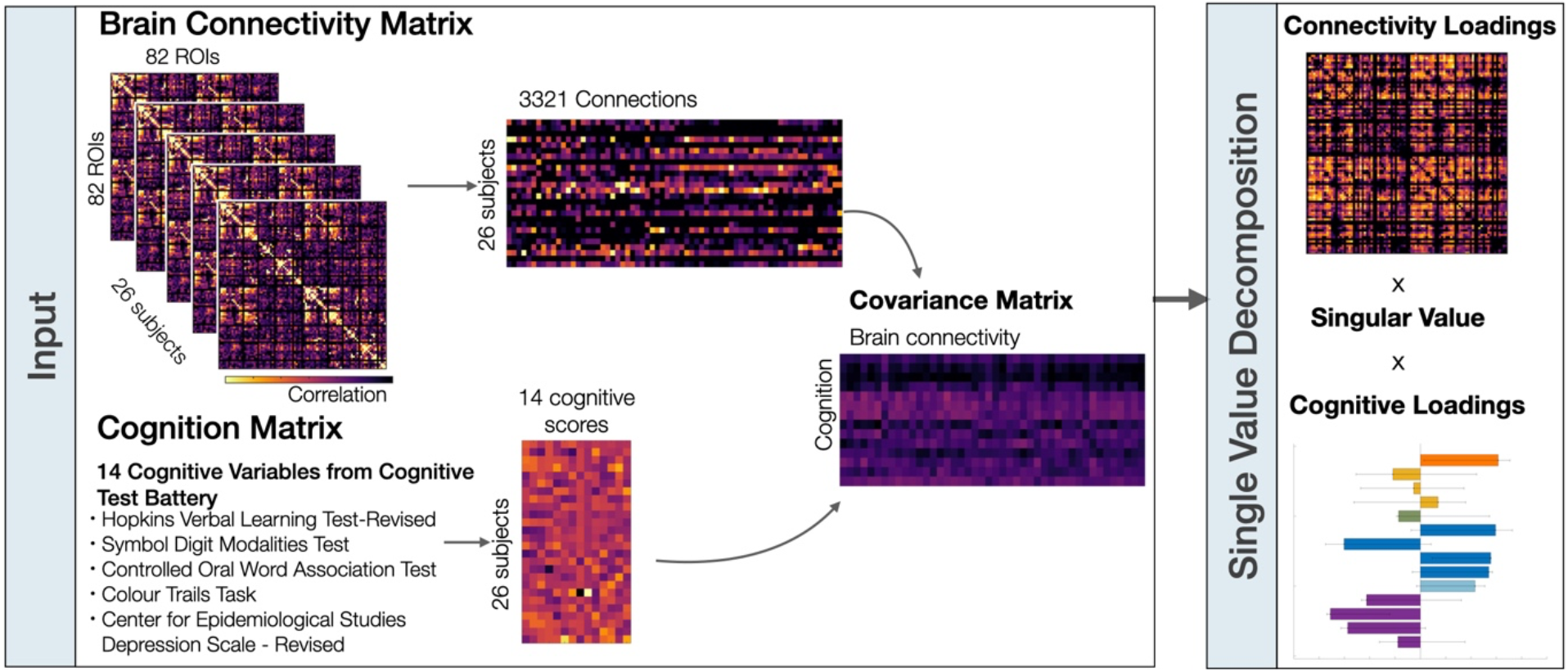
Overview of partial least squares analyses. Two partial least squares analyses were performed on both brain connectivity data sets (i.e., hemodynamic connectivity, metabolic connectivity) separately. The brain connectivity matrices were first sorted by stacking the upper triangle elements from each participants’ matrices. The rows of the brain connectivity and cognitive matrices correspond to participants and the columns correspond to either the brain connections or cognitive scores. The covariance between the brain connectivity and cognition matrices was computed across participants, resulting in a rectangular connectivity-cognition covariance matrix. This covariance matrix was then subjected to singular value decomposition. Refer to Materials and Methods for details.

## Results

We first provide an overview of the cognitive outcome variables of the neuropsychological test battery. Next, we describe the hemodynamic (i.e., fMRI functional connectivity) and metabolic connectivity (i.e., FDG-fPET functional connectivity) across participants. Finally, we show how both connectivity maps relate to cognition and quantify their differences.

### Cognitive measures

Participants completed a neuropsychological test battery that described distinct cognitive domains across 14 outcome variables (Table 1). Most cognitive variables correlated significantly within each cognitive test, but not across tests (Figure 2) suggesting each cognitive test measured distinct cognitive domains. An exception was that individuals with higher depression scores on the CESD-R were overall slower during congruent trials of the Stroop task (i.e., reading colour names; r(24) = 0.54, 95% CI [0.28, 0.83], p < 0.05). Also, performance during the Symbol Digits Modality test, correlated negatively with performance during the second part of the Colour Trail test (CT2 score) (r(24) = -0.48, 95% CI [-0.83, - 0.28], p < 0.05). For the Partial Least Squares analyses, the total score from the Stroop congruent trials was removed as there was no variability across participants, as all participants received the maximum score of 112.

**Table 1.**
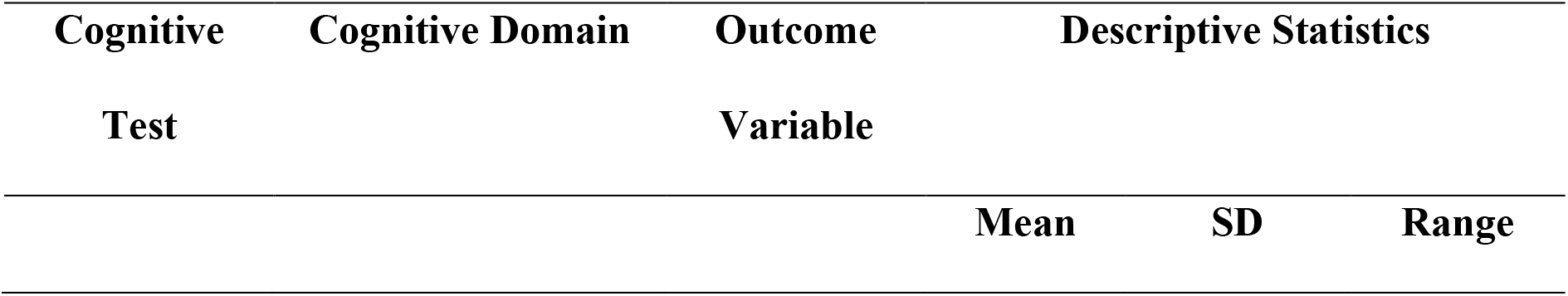
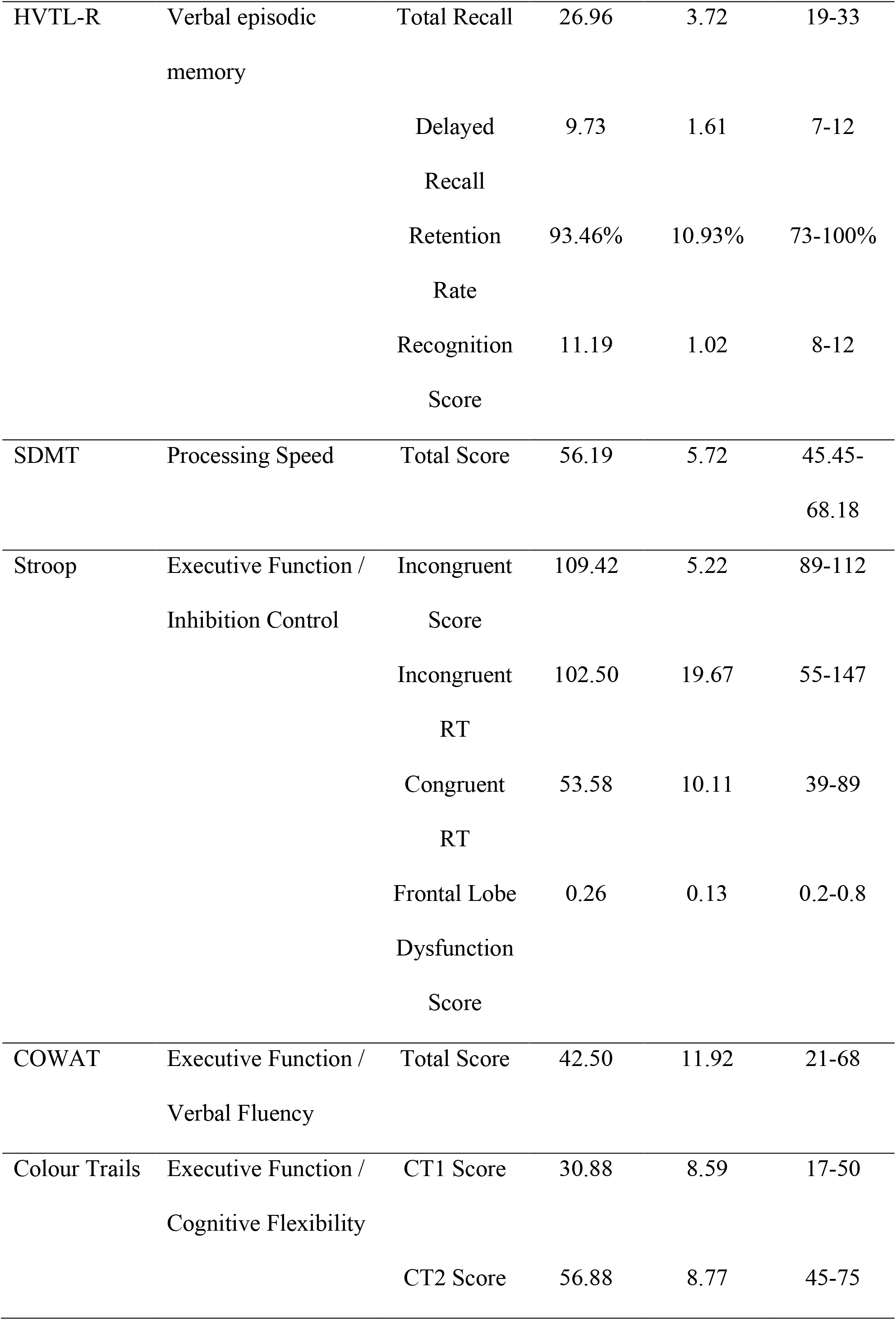
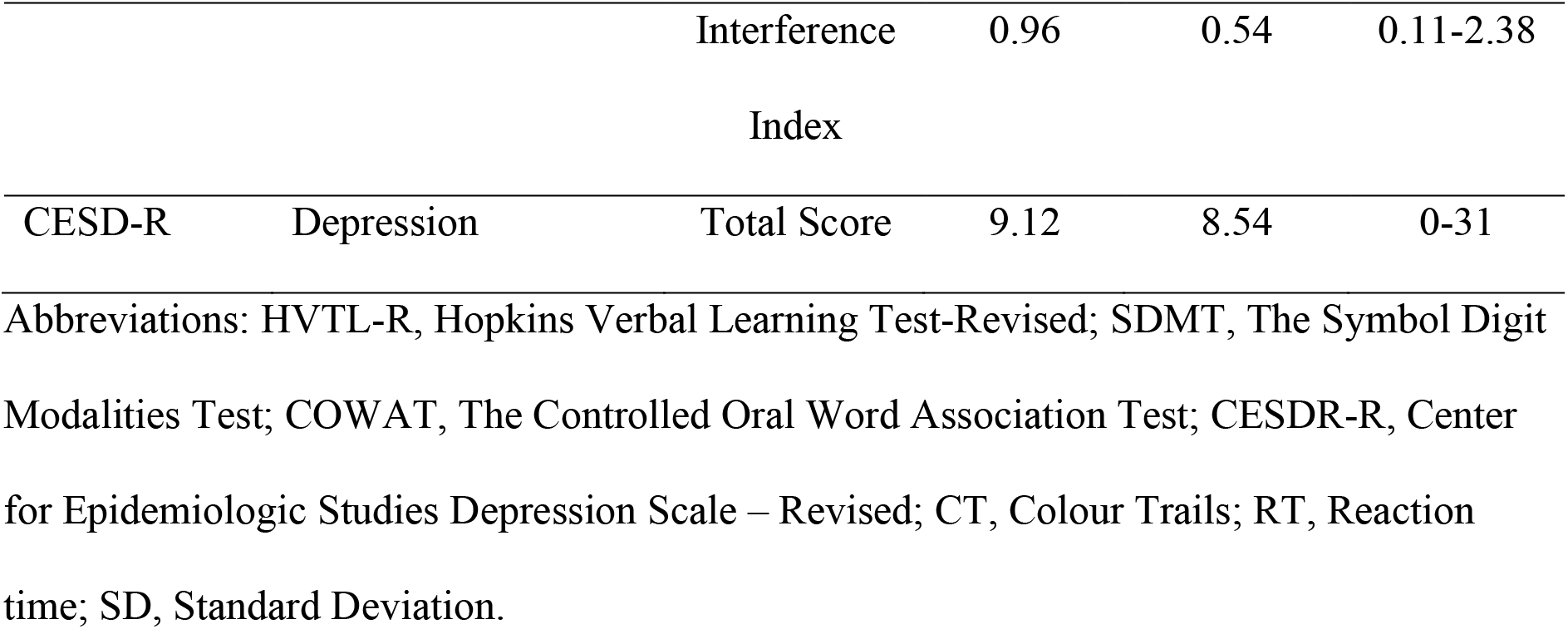
Cognitive outcome variables from the cognitive battery.

**Figure 2.**
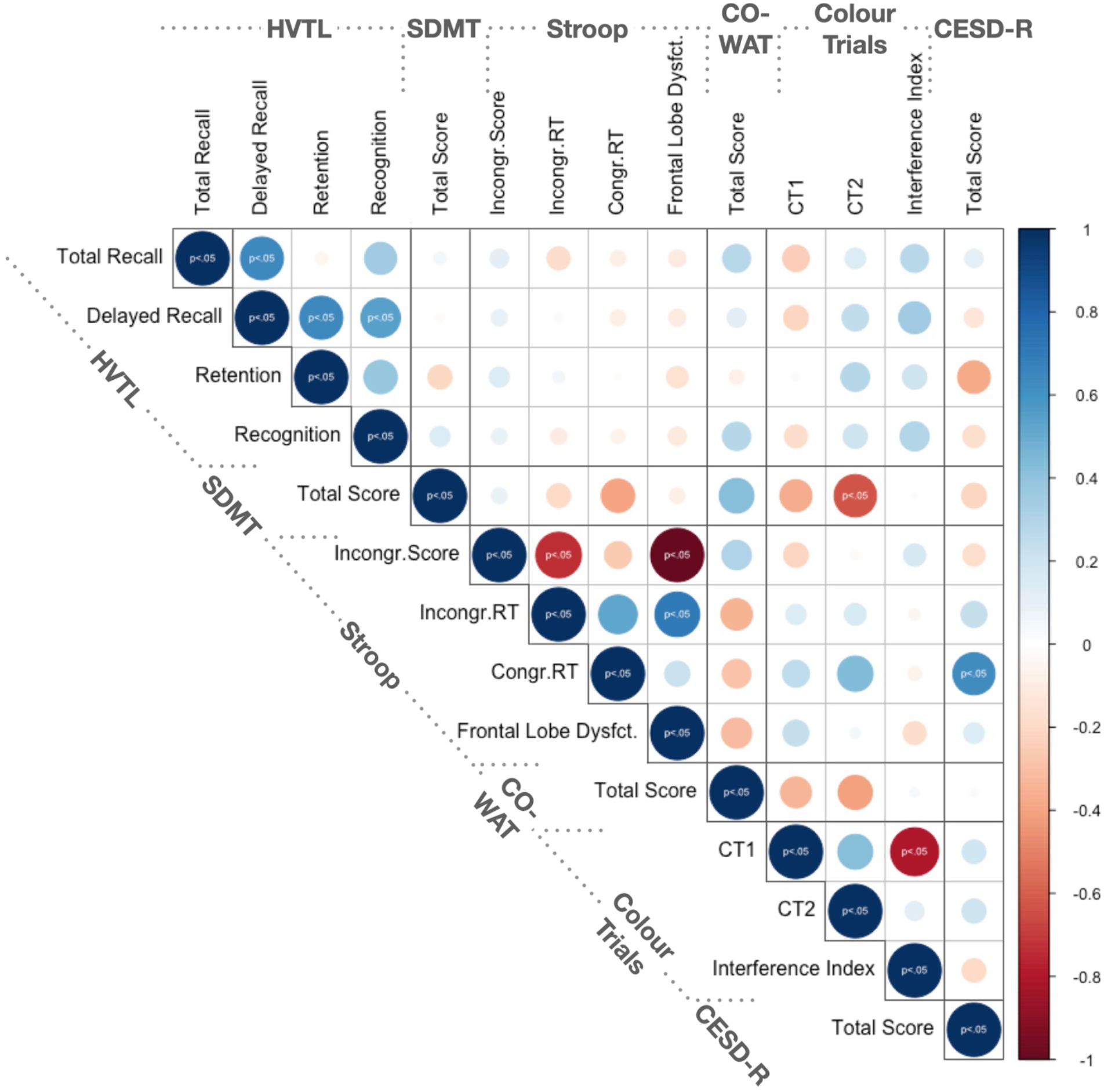
Correlation matrix of 14 cognitive outcome variables obtained from the neuropsychological test battery. Significant relationships are indicated with p > 0.05 corrected for multiple comparisons using false discovery rate (Benjamini and Hochberg, 1995). Pearson’s correlation was performed for continuous and Spearman correlation was performed for ordinal data. Positive relationships (0 ≤ r ≤ 1) are indicated in blue and negative relationships (0 > r ≤ -1) are indicated in red. Circle size corresponds to the absolute size of the correlation coefficient as indicated by the blue-red coloured scale. Abbreviations: HVLT, Hopkins Verbal Learning Test-Revised; SDMT, Symbol Digit Modality Test; COWAT, Controlled Oral Word Association Test; CESD-R, Center for Epidemiologic Studies Depression Scale – Revised; CT1, Colour Trails 1; CT2, Colour Trails 2; RT, Reaction Time.

### Hemodynamic and Metabolic Connectivity

The haemodynamic and metabolic connectomes have been reported previously (Jamadar et al., 2021; Jamadar et al., 2020) and are included here for completeness. The hemodynamic connectome (Figure 3a) showed medium to strong connectivity within most anatomical subdivisions, both within and between hemispheres. The strongest hemodynamic connectivity (r ≥ 0.7) was found bilaterally in the frontal, parietal, and occipital anatomical subdivisions. A number of strong long-range connections included frontoparietal, parieto-occipital, and temporoparietal regional connectivity. These long-range connections were evident both within and between hemispheres but were of smaller magnitude than the short-range and homotopic connections. Subcortical and orbitofrontal regions were the least interconnected regions in the BOLD-fMRI data (r ≥ 0.2).

**Figure 3.**
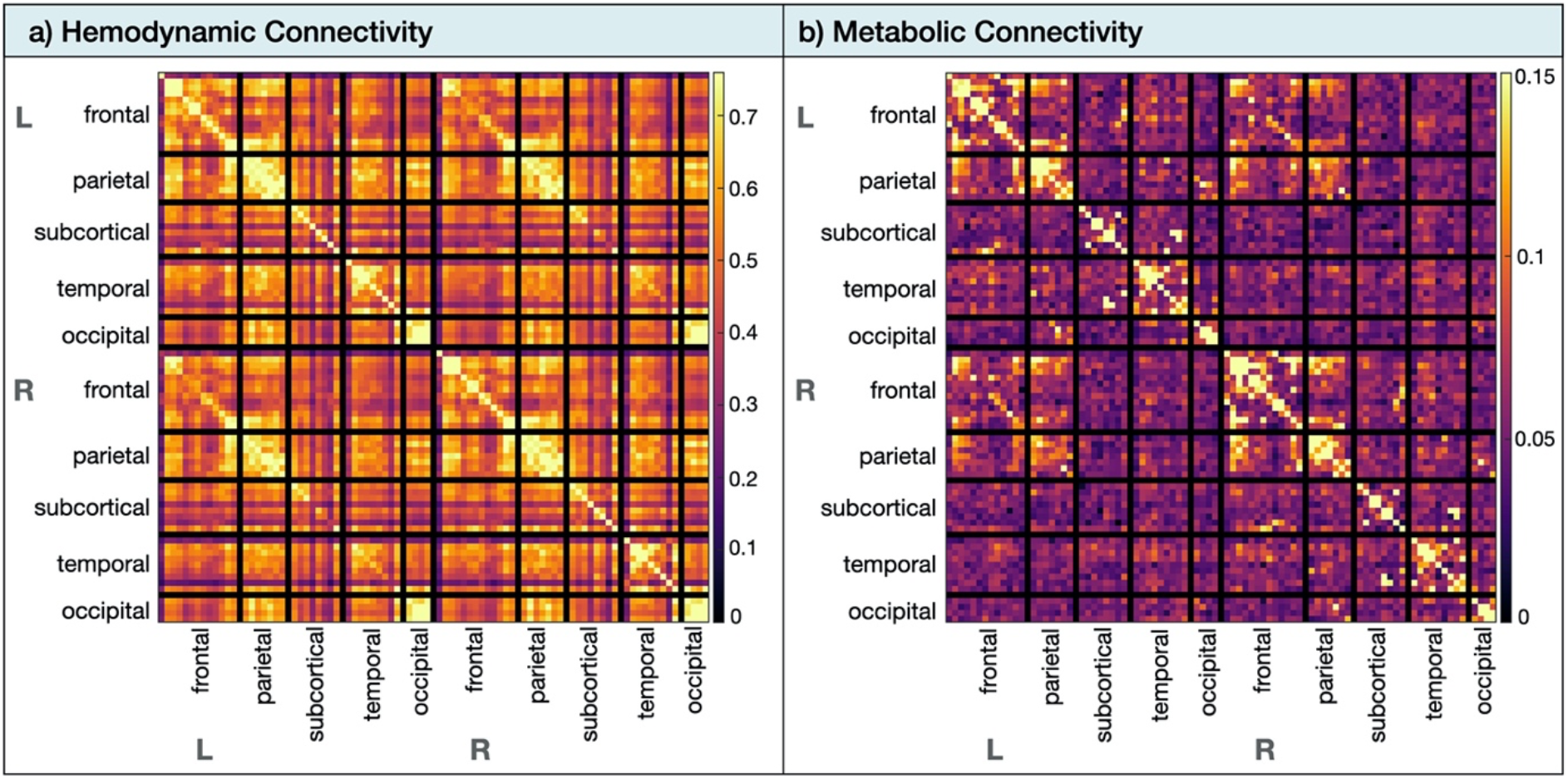
Hemodynamic and metabolic connectivity at rest. (a) The hemodynamic (i.e., fMRI functional connectivity) was thresholded from 0 < r < 0.76. (b) The metabolic connectivity (i.e., fPET functional connectivity) was thresholded from 0 < 0 < 0.15. Abbreviations: L, left; R, right.

The metabolic connectome (Figure 3b) showed the strongest connectivity (r ≥ 0.15) within the frontoparietal areas, which was more apparent within than between hemispheres. Left–right homotopic connectivity was not visually apparent for subcortical, temporal, and occipital cortices.

### Partial Least Squares Results

The Partial Least Squares (PLS) analyses applied to the fMRI and fPET data sets separately identified one significant latent variable that described the relationship between the hemodynamic connectivity and cognition, and one significant latent variable that described the relationship between the metabolic connectivity and cognition.

#### Hemodynamic Connectivity and Cognition Relationship

The PLS analyses revealed that one latent variable captures the relationship between hemodynamic connectivity pattern and cognition (67.13 % of total covariance; singular value = 39.43, p = 0.003, permutation testing with 5,000 iterations). The distribution of cognitive loadings revealed that each cognitive variable within each test (Table 1) in general loaded uniformly in their direction onto the latent variable (Figure 4ai). For example, all sub-scales of the HVLT loaded negatively onto the latent variable, and all sub-scales of the Stroop loaded positively. Bootstrapped confidence intervals revealed that three cognitive variables were expressed the strongest by the latent variable: participant’s depression score (CESD-R score; loading = 0.61, 95% bootstrapped CI [0.15,0.82]), inhibition control speed (i.e. the response time in naming a font colour of an incongruent word during the Stroop task; loading = 0.56, 95% bootstrapped CI [0.18,0.57]), and memory retention (HVTL retention score; loading = -0.71, 95% bootstrapped CI [-0.34, -0.72). The hemodynamic connections all loaded strongly positively onto the latent variable (loading > 0.67; Figure 4aii). The strongest loadings (r ≥ 0.85) were found bilaterally in the frontal and parietal anatomical subdivisions. Thresholding the connectivity matrix at the 99^th^ percentile (Jamadar et al., 2020), revealed that 41.4% of the strongest connections were part of the frontal cortex (e.g., superiorfrontal, middlefrontal, parstriangularis, parsopercularis, medialorbitofrontal, precentral and rostralanteriorcingulate), and 24.1% of the total strongest connections were part of the parietal cortex (e.g., supramarginal, posteriorcingulate, precuneus, isthmuscingulate). The subcortical areas contained (e.g., caudate, hippocampus, insula, putamen) and the temporal cortex contained 17.2% of the total strongest connections (i.e., superiortemporal, fusiform, banks), respectively. There were no strong connections in the occipital cortex. Interpreting the cognition loadings together with the brain loadings, the PLS analysis revealed that higher depression, higher inhibitory control speed and lower memory retention are associated with higher hemodynamic connectivity particularly in the frontal and parietal anatomical interhemispheric subdivisions.

**Figure 4.**
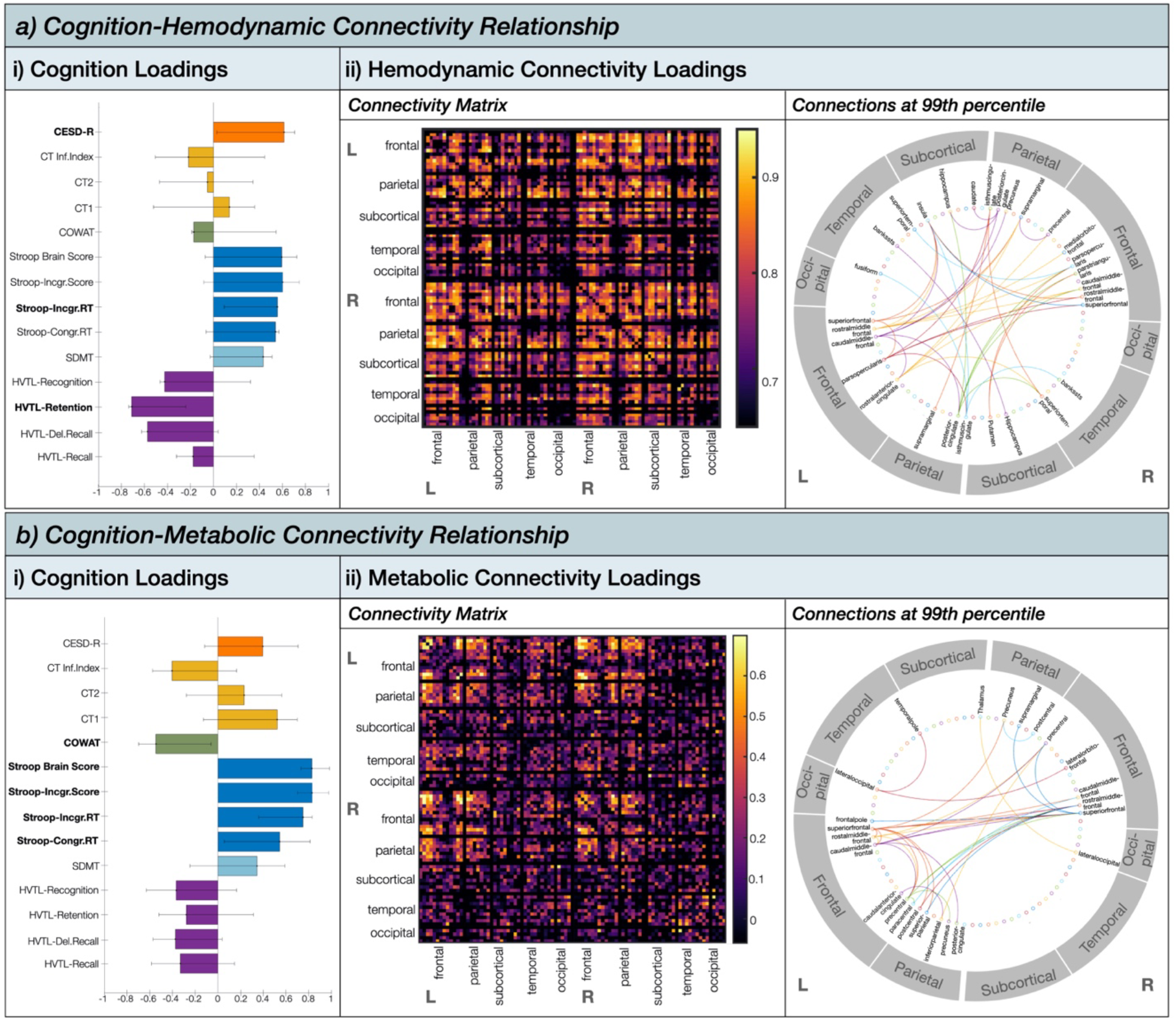
Partial least squares analysis results showing the a) cognition-hemodynamic connectivity relationship, and b) cognition-metabolic connectivity relationship. (i) Cognition loadings for significant latent variable. Error bars represent 95% Confidence intervals from bootstrap resampling (5,000 iterations). (ii) Connectivity loadings for significant latent variable. The circular plot shows the strongest anatomical connections thresholded at the 99^th^ percentile.

#### Metabolic Connectivity and Cognition Relationship

The PLS analyses revealed that one latent variable captures the relationship between metabolic connectivity pattern and cognition (30.77 % of total covariance; singular value = 24.42, p = 0.04, permutation testing with 5,000 iterations). The distribution of cognitive loadings revealed that each cognitive variable within each test (Table 1) loaded generally uniformly and in their direction onto the latent variable (Figure 4bi). The direction of the cognitive loadings was similarly expressed by the latent variable describing the hemodynamic connectivity-cognition relationship. Bootstrapped confidence intervals revealed that all outcome variables of the Stroop task, measuring executive functioning/inhibitory control, loaded strongly positively onto the latent variable. Further, the COWAT score loading = -0.58, 95% bootstrapped CI [-0.15, -0.79]), measuring executive functioning/verbal fluency was also expressed strongly negatively by the latent variable. The metabolic connections loaded mostly positively (loading > 0.54) onto the latent variable (Figure 3bii). However, there were also a few connections that loaded negatively, although very weakly (loading < -0.15) (Appendix Figure S2). These negative loadings were distributed across the brain. The strongest loadings (r ≥ 0.54) all loaded positively and were found predominantly in the frontal and parietal anatomical subdivisions. Thresholding the connectivity matrix at the 99^th^ percentile, revealed that 50% of the strongest connections were part of the frontal cortex (e.g., frontal pole, superior frontal, middle frontal, lateral orbitofrontal, caudal anterior cingulate, precentral) and 33% of the total strongest connections were part of the parietal cortex (e.g., postcentral, posterior cingulate, pre-cuneus, inferior parietal, superior parietal, supra-marginal). The occipital cortex contained only 8.3% of the total strongest connections (i.e., lateral occipital) and the subcortical (i.e., thalamus) and temporal cortex (i.e., temporal pole) only 4.2%, respectively. Interpreting the cognition loadings together with the brain loadings, the PLS analysis revealed that higher inhibitory control and lower verbal fluency are associated with predominantly higher metabolic connectivity particularly in the frontal and parietal anatomical subdivisions.

#### Differences in Hemodynamic-Cognition and Metabolic-Cognition Relationship

To compare the connections that contributed to the cognition-metabolic connectivity relationship and those that contributed to the cognition-hemodynamic relationship, we computed the scalar dot product between the brain saliences (U) of the significant latent variable from both PLS analyses. A cosine value of 1 means that the saliences are identical and 0 means orthogonality or no correlation. This analysis revealed a cosine similarity of 0.23 (i.e., weak relationship) indicating that the effects of the PLS for the hemodynamic-cognition relationship differed from the effects from the metabolic-cognition relationship. This was confirmed by the similarity matrix of each relationship’s brain loadings, showing overall little overlap across the two modalities with the most correlation coefficients ranging between -0.1 and 0.1 (Figure 5). The highest similarity between the two modalities (r > 0.4) was evident for the frontal and parietal cortex for both hemispheres. The loading matrices were anticorrelated (loading < -0.3) for occipital and temporal subdivisions.

**Figure 5.**
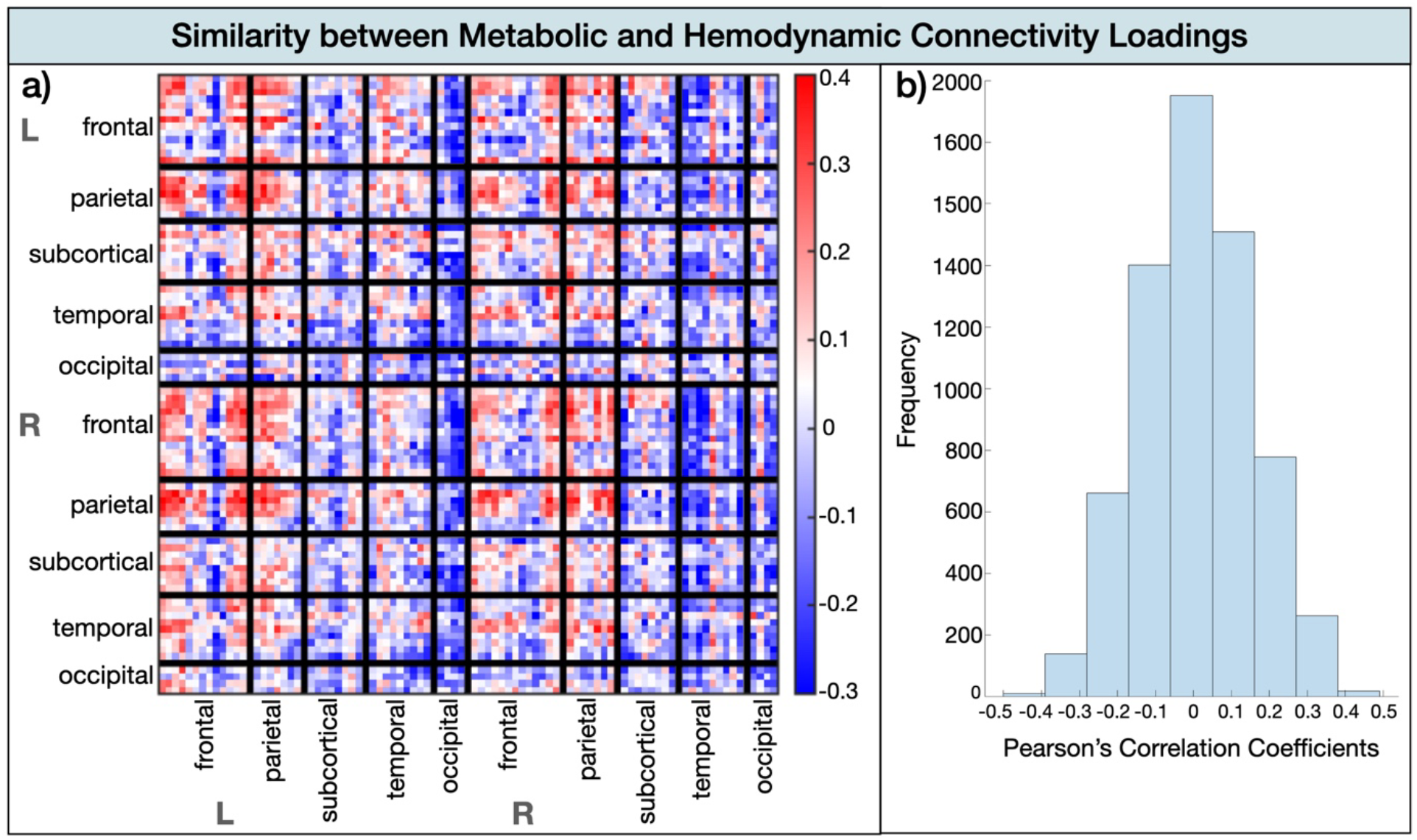
Similarity between the metabolic and hemodynamic connectivity loadings. (a) Similarity matrix by brain area. (b) Histogram of Pearson’s correlation coefficients indicating the frequency of similarity strength.

## Discussion

The present study used simultaneous resting-state FDG-PET/fMRI to investigate, for the first time, how spatially distant synchronous brain signals measured via cerebrovascular hemodynamic responses (i.e., fMRI; hemodynamic connectivity) and glucose uptake (i.e., FDG-PET; metabolic connectivity) relate to a range of cognitive functions. Our simultaneous fPET and fMRI acquisition at a high temporal resolution enabled multimodal within-subject analyses of resting-state brain activity without the confound of intra-individual differences (e.g., fatigue, nutrient intake, blood chemistry) that occur when measuring both modalities not simultaneously. We applied Partial Least Squares (Krishnan et al., 2011; McIntosh and Lobaugh, 2004) to extract latent variables capturing the maximum covariance between hemodynamic and metabolic connectivity matrices with 14 cognitive measures, including episodic memory, processing speed, executive functioning, and depression. Results revealed that one latent variable captured the relationship between hemodynamic connectivity and cognition and one latent variable captured the relationship between metabolic connectivity and cognition. The cognitive battery was indexing orthogonal cognitive domains. This supports the global connectome-cognition view, which states that a global cognitive factor is accounted by a single set of connections (Goyal et al., 2020b; Smith et al., 2015b). By contrast, our results do not support the domain-specific connectome-cognition view, which would suggest that distinct sets of connections are required to support cognition (e.g., Ziegler et al., 2013; Zimmermann et al., 2018).

Although cognition was expressed globally by one set of connectivity-cognition latent variable, the specificity of how hemodynamic and metabolic connectivity related to cognition varied. For both modalities the frontoparietal anatomical subdivisions related the strongest to cognition (Figure 4), however for hemodynamic responses this network expressed executive functioning, episodic memory, and depression, whereas for metabolic responses this network exclusively expressed executive functioning. This is compatible with the argument that metabolic and hemodynamic connectivity provide unique, but complementary insights into cognition (Chen et al., 2018; Hahn et al., 2020; Jamadar et al., 2021; Mier and Mier, 2015; Sala and Perani, 2019; Yakushev et al., 2017).

### A global set of metabolic and hemodynamic connections map onto cognition

Our results support the contention that the overall wiring of a connectivity network has a domain-general role in cognition. Critically, this domain-general characteristic is shared by both the metabolic and hemodynamic processes, indicating that it is a shared characteristic across multiple physiological levels of the human connectome. This finding is in line with classical theoretical proposals that brain networks exhibit a flexible architecture with their functional network assignment to adaptively process changing cognitive demands (Dehaene et al., 1998; Duncan, 2001; Miller and Cohen, 2001). Flexible, domain-general interactions likely allow different information to become quickly integrated and exchanged, leading to a dominant pattern of co-activation across different cognitive states.

In our results, the frontoparietal anatomical subdivisions emerged as the dominant regions supporting a domain-general role in cognition. The frontoparietal anatomical network was previously coined a multi-demand system that is co-activated when performing a diverse range of cognitive demanding tasks, including selective attention, working memory, task switching, response inhibition, conflict monitoring, learning or problem solving (Assem et al., 2020; Chein & Schneider, 2005; Cole et al., 2014; reviewed by Marek & Dosenbach, 2018). In line with this general systems role to support information integration and exchange that mediates cognitive operations, damage to the frontoparietal network has been reported to be associated with disorganised behaviour and decreased fluid intelligence (Hearne et al., 2016). Further, this system has been shown to play domain-general protective role against mental health symptoms such as depression (Schultz et al., 2018).

### Metabolic and functional connectivity relate to distinct aspects of cognition

The behavioural variables loaded uniformly on the latent variables for the metabolic connectivity-cognition and hemodynamic connectivity-cognition pattern but differed in their loading strengths. The fPET metabolic and BOLD-fMRI hemodynamic connectivity had the strongest network configuration in frontoparietal cortices. However, this network seems to relate to distinct cognitive functions for both imaging modalities. Specifically, the resting-state hemodynamic connectivity in this network was positively associated with inhibition, depression and negatively with memory retention. The resting-state metabolic connectivity in this network in turn was associated positively with executive functioning and inhibition; and negatively with executive functioning and verbal fluency.

The cognition-connectivity pattern revealed by fMRI is in strong accord with numerous previous fMRI studies revealing the brain mechanisms underlying cognition. For example, the frontoparietal network, particularly involving the anterior cingulate cortex, precuneus or posterior cingulate cortex, has been shown to be a core network involved in cognitive control monitoring and the facilitation of conflict resolution during a task (Botvinick et al., 2004; Shenhav et al., 2013). Additionally, this flexible and domain-general hub has also been involved in emotional processing, clinical symptoms such as depression (Schultz et al., 2018), and memory (Wallis et al., 2015). These findings are corroborated in the cognition-connectivity patterns observed in this study. In addition to frontoparietal co-activation, the hemodynamic connectivity loadings were also prevalent in cortico-cortical networks, for example involving the insula or hippocampus. The insula is strongly interconnected with frontal and parietal areas supporting its role as a major multimodal network hub that underpins cognition, memory and emotional processing (Contreras et al., 2012; Menon and Uddin, 2010). The hippocampus supports a vast array of memory functions, such as retaining information across delays (Jeneson et al., 2011; Müller et al., 2018).

In contrast to the hemodynamic connectivity-cognition relationship, the latent variable expressing the metabolic connectivity-cognition relationship was strongly localised in the frontoparietal areas and associated exclusively with executive functioning. Previous studies have reported that resting-state metabolic connectivity is particularly evident in frontoparietal areas (Shokri-Kojori et al., 2019; Yakushev et al., 2017; Hahn et al., 2019). Here we extend these finding by observing that the co-activation at rest is behaviourally relevant in supporting executive control. We note the existence of a small proportion of negative connections (only 25.22 % of connections) that contributed to the cognition-metabolic connectivity relationship. These negative cognition-connectivity associations can reflect either reduced positive associations or anti-correlations (Hearne et al., 2016). There is also the possibility that these scattered negative loadings (Figure S2) might be a pre-processing epiphenomenon (Jamadar et al., 2020). Future research is needed to investigate whether the small fraction of negative associations in the metabolic connectome are behaviourally meaningful.

The apparent specificity of the cognition-metabolic connectivity relationship, i.e., the exclusive focus on frontoparietal cortices, may be indicative of signal artefacts in either the FDG-fPET or BOLD-fMRI, i.e., reduced signal-to-noise or non-neuronal confounders, respectively. The reduced sensitivity of the FDG-fPET signal must be noted as the processing pipeline, including filters and models, are immature compared to the years of advanced development that has been dedicated to BOLD-fMRI signal processing as reported in the scientific literature. Conversely, this advancement has potentially led to the identification of non-neuronal confounders and spatial artefacts in BOLD-fMRI that are not present in the FDG-fPET signal, such as magnetic field and haemoglobin-based artefacts (Liu, 2017; Ward et al., 2020). The disparity in the results from the two modalities is augurs well for gaining deeper insights to improve our understanding of cognition-brain connectivity relationships.

In conclusion, this study is an important step in revealing that cognition is supported by a domain-general hemodynamic and metabolic processing. Crucially, the metabolic processes appear to be more spatially defined by frontoparietal areas, whereas the hemodynamic processes throughout the frontal, parietal, temporal, and occipital areas collectively support cognition. These findings demonstrate the unique advantages that simultaneous FDG-PET/fMRI has to provide a comprehensive understanding of the neural mechanisms that underpin cognition, and highlights the importance of multimodality imaging in cognitive neuroscience research.

## Materials and Methods

All methods were reviewed by the Monash University Human Research Ethics Committee, following the Australian National Statement of Ethical Conduct in Human Research (2007). Participants provided informed consent to participate in the study. Administration of ionising radiation was approved by the Monash Health Principal Medical Physicist, following the Australian Radiation Protection and Nuclear Safety Agency Code of Practice (2005). Data from this study is available on OpenNeuro with the accession number ds002898. The Data Descriptor for this study with detailed acquisition and validation analyses is provided in Jamadar et al. (2020), and results of the comparison between fPET, static PET, and BOLD-fMRI connectomes is presented in Jamadar et al. (2021).

### Participants

Twenty-eight participants were recruited from the general community. An initial screening interview assessed that these participants had no history of hypertension or diabetes, had no neurological and psychiatric illness, or were on psychoactive medication affecting cognitive functioning or cerebral blood flow. Participants were also screened for claustrophobia, non-MR compatible implants, clinical or research PET scan in the past 12months, and women were screened for current or suspected pregnancy. Prior to the scan, participants were directed to consume a high-protein/low-sugar diet for 24h, fast for 6h, and drink 2-6 glasses of water. Blood sugar level was measured using an Accu-Check Performa (model NC, Mannheim, Germany); all participants had blood sugar levels <10mmol/L with none exceeding 4.73 mmol/L. Two participants were excluded for further analyses, as one participant did not complete the full scan and the infusion pump failed for one participant. The total sample (n = 26, 77% females) were aged between 18-23 years (mean age = 19.50 years, SD = 1.36 years), right-handed (Oldfield, 1971), English speakers (Table S1 for summary demographics). Although the sample consisted of significantly larger proportion of females, there were no significant (p>0.5) gender-based differences observed in their demographics (Figure S1).

### Neuropsychological Test Battery

Prior to the scan, participants completed a test battery consisting of six neuropsychological test or scales assessing a wide range of cognitive functioning: (1) Hopkins Verbal Learning Test-Revised (Benedict et al., 1998), (2) Symbol digit modalities test (Smith, 1991), (3) Stroop Neuropsychological Screening test (Trenerry et al., 1989), (4) single-letter controlled oral word association test (COWAT; Ruff et al., 1996), (5) Colour Trails Task (Reitan, 1958), and the (6) Center of Epidemiologic Studies Depression Scale – Revised (Radloff, 1977). Full details of the neuropsychological tests are provided in the Appendix.

Overall, the five tests produced 14 cognitive outcome variables, which are summarised in Table 1 (Results). Relationships between the cognitive outcome variables were explored via Pearson’s correlations for continuous outcome variables and Spearman correlation for ordinal outcome variables. Relationships were considered significant at a false discovery rate corrected p value of 0.0062 (Benjamini and Hochberg, 1995)

### Simultaneous MR-PET data acquisition

Following the completion of the cognitive battery (approximately 30 minutes), participants underwent preparation for the simultaneous MR-PET scan. They were first cannulated in the vein in each forearm, and a 10-ml baseline blood sample was taken. For all participants, the left cannula was used for FDG infusion, and the right cannula was used for blood sampling.

Participants underwent a 95-min simultaneous MR-PET scan in a Siemens (Erlangen) Biograph 3-Tesla molecular MR scanner. Participants were positioned supine in the scanner bore with their head in a 16-channel radiofrequency head coil and were instructed to lie as still as possible with eyes open and think of nothing in particular. FDG (average dose 233 MBq) was infused over the course of the scan at a rate of 36 mL/h using a BodyGuard 323 MR-compatible infusion pump (Caesarea Medical Electronics, Caesarea, Israel). Infusion onset was locked to the onset of the PET scan.

Plasma radioactivity levels were measured throughout the duration of the scan. At 10-minutes post-infusion onset, a 10 mL of blood sample was taken from the right forearm using a vacutainer; the time of the 5-mL mark was noted for subsequent decay correction. Subsequent blood samples were taken at 10-minute intervals for a total of 10 samples for the duration of the scan. Immediately following blood sampling, the sample was placed in a Heraeus Megafuge 16 centrifuge (ThermoFisher Scientific, Osterode, Germany) and spun at 2000 rpm for 5 minutes; 1000 μL plasma was pipetted, transferred to a counting tube, and placed in a well counter for 4 minutes. The count start time, total number of counts, and counts per minute were recorded for each sample. The average radioactivity concentration persistently increased over time with the lowest relative rate occurring at the end of the acquisition.

### MRI pre-processing

For the structural T_1_ image, the brain was extracted in Freesurfer, then registered to MNI152 space using Advanced Normalization Tools (ANTs). The gray matter, white matter, and brain cortex labels of the structural T_1_ image were segmented into 82 regions using Freesurfer with Desikan-Killiany Atlas (Diedrichsen et al., 2009).

The six blocks of EPI scans for all participants (a total of 1452 volumes) underwent a standard fMRI pre-processing pipeline. Specifically, all scans were brain extracted (FSL BET, Smith, 2002), motion corrected (FSL MCFLIRT, Jenkinson et al., 2002), slice timing corrected (FSL, using Fourier-space time-series phase-shifting) and band-pass filtered (0.1>Hz > 0.01) to remove low-frequency noise (FSL, Jenkinson et al., 2012), and spatially smoothed using a Gaussian kernel of FWHM of 8mm. Across subjects, the average mean framewise translation motion was 0.41 mm, maximum was 1.09 mm.

### PET image reconstruction and pre-processing

The 5700-s list-mode PET data for each subject were binned into 356 3D sinogram frames each of 16-s interval. The attenuation for all required data was corrected via the pseudo-CT method (Burgos et al., 2014). Ordinary Poisson-Ordered Subset Expectation Maximization algorithm (3 iterations, 21 subsets) with point spread function correction was used to reconstruct 3D volumes from the sinogram frames. The reconstructed DICOM slices were converted to NIFTI format with size 344×344×127 (voxel size: 2.09×2.09×2.03 mm ^3^) for each volume. A 5-mm FWHM Gaussian postfilter was applied to each 3D volume. All 3D volumes were temporally concatenated to form a 4D (344 × 344 × 127 × 356) NIFTI volume. A guided motion correction method using simultaneously acquired MRI was applied to correct the motion during the PET scan. We retained the 225 16-s volumes commencing from the 30-minute timepoint, which matched the start of the BOLD-fMRI EPI acquisition, for further analyses.

The 225 PET volumes were motion corrected (FSL MCFLIRT, Jenkinson et al., 2002); the mean PET image was brain extracted and used to mask the 4D data. The fPET data were further processed using a spatiotemporal gradient filter to estimate the short-term change in glucose uptake from the cumulative glucose uptake that was measured (Jamadar et al., 2020). The filter removed the accumulating effect of the radiotracer and other low-frequency components of the signal to isolate short-term resting-state fluctuations. This approach intrinsically adjusted for the mean signal while avoiding global-signal regression and other approaches that may create spurious anticorrelations in the data (Murphy and Fox, 2017). Due to radiotracer dynamics, it was not expected that the fPET sensitivity would be uniform across the 60 minutes of the resting-state data acquisition. As the radiotracer accumulated in the brain, it was anticipated that the signal-to-noise ratio (SNR) of the PET image reconstruction would progressively improve. The spatiotemporal filter has been described extensively in our previous work (Jamadar et al., 2021; Jamadar et al., 2020).

### Functional and Metabolic Connectivity Analyses

For fPET and fMRI, time series were extracted for each of the 82 regions of interest (ROIs) from the segmentation of the T1 - weighted image, interpolated using an ANTs rigid registration (Avants et al., 2011). To construct a connectivity matrix, Pearson’s correlation coefficients were estimated between the timeseries from pairs of regions. This produced a per-subject per-modality 26 × 82 × 82 matrix corresponding to the 60 minutes of resting-state in the experimental protocol. For interpretation of connectivity patterns, the 82 ROIs were anatomically sorted as defined by the Desikan-Killiany Atlas (i.e., frontal, parietal, occipital, subcortical, temporal; Diedrichsen et al., 2009).

### Partial Least Squares Analyses

We used Partial Least Squares (PLS) analyses to assess the multidimensional functional relationships between (1) the hemodynamic connectome and cognition, as well as (2) the metabolic connectome and cognition (Krishnan et al., 2011; McIntosh and Lobaugh, 2004) (Figure 1). PLS is an unsupervised multivariate machine learning technique that extracts the common information between two datasets (i.e., brain connectivity [**X**] and cognitive responses [**Y**]) by finding orthogonal sets of latent variables with maximum covariance, which reflect the linear combinations of the original data. In our case, the brain connectivity is either the hemodynamic connectome [**X**_**F**_] or metabolic connectome [**X**_**M**_]. Prior to the application of PLS, the upper triangle of the hemodynamic connectivity matrix and the metabolic connectivity matrix (i.e., 3321 connections, respectively) were vectorized and stacked as participant by connection resulting in matrices sized 26 × 3321, respectively. The cognition matrix was sized at 26 × 14. These subject-specific the hemodynamic connectivity matrix (**X**_**F**_), metabolic connectivity (**X**_**M**_) matrices and the cognitive response matrix (**Y**) were subsequently z-scored column-wise.

First, the correlation matrix between the brain connectivity (**X**) and cognition matrix (**Y**) is computed **R** = **X**^**T**^**Y** and single value decomposition (Eckart and Young 1936) is next applied to that connectivity-cognition matrix, resulting in **R = USV**^**T**^. The outcome of the single value decomposition is a set of mutually orthogonal latent variables, whereby **U** and **V** are orthogonal matrices consisting of left and right singular vectors and **S** is a diagonal matrix of singular values. The number of latent variables is equal to the rank of the covariance matrix R, which is the smaller of its dimensions. Every latent variable is associated with 1) a singular value (diagonal elements of **S**) indicated the correlation explained by that latent variable, 2) a vector of singular values **U**, which represent the behavioural saliences, and 3) a vector of singular values **V**, which represent the brain saliences. The behavioural saliences indicate how strong each one of the cognitive variables contributes to the brain-design correlation explained by a particular latent variable. Similarly, the brain saliences **V** express how strong every connection contributes to the brain-design correlation explained by a particular latent variable. The projection of every subject’s original connectivity (in **X**) onto the multivariate brain salience pattern (in **V**) results in brain scores **L**_**x**_ = **XV**. Brain scores measure the similarity of a subject’s individual brain data with the salient brain pattern. Similarly, cognitive scores can be computed by **Ly** = **YU**, which represent a projection of every subject’s design variable onto the respective design saliences. Finally, brain loadings (or weights) were computed as the Pearson’s correlation between the brain connectivity matrix and the PLS analysis-derived brain scores. Similarly, cognitive loadings were computed as the Pearson’s correlations between cognitive variables and the PLS analysis-derived cognition scores across the cohort. Loadings can be interpreted as indexing the degree contribution of each variable to the PLS analysis-derived latent variable. Loadings are only interpreted for significant latent variables.

The significance of each latent variable is assessed via permutation tests (5000 iterations) of the singular values from the single value decomposition of the brain and cognition matrices and the reliability of each connectivity estimate to the latent variable is assessed via bootstrap resampling (5000 iterations). The reliability of the loading of each connection onto the brain-cognition relationship in each latent variable is established via bootstrap (5000 iterations). A connection with a positive bootstrapped loading contributes positively and reliably to the brain-cognition correlation obtained for that latent variable, whereas a connection with a negative high bootstrapped loading contributes negatively and reliably to the brain-cognition relationship. Bootstrapping is also used to construct 95% confidence intervals on the brain-cognition correlations.

### Hemodynamic versus metabolic connectivity in relation to cognition

To compare the brain connections that contributed to the hemodynamic connectome-cognition relationship and metabolic connectome-cognition relationship, the scalar product between the brain saliences (**U**) resulting from each PLS were computed for significant latent variables. Similarly, to compare the cognitive responses that contributed to the metabolic connectome-cognition relationship and functional connectome-relationship, we calculated the dot product between the behavioural saliences (**V**) that resulted from both PLS analyses. A scalar product of 0 suggests no overlap across modalities and a scalar product of 1 suggest strong overlap across modalities (i.e., fPET and fMRI). Finally, to identify the anatomical location of similar brain loadings across modalities, Pearson’s correlations was performed on the brain loadings matrices of both modalities. This results in a matrix of cosine similarity between the two modalities.

## Acknowledgements

This work was supported by an Australian Research Council (ARC) Linkage Project (LP170100494) that includes financial support from Siemens Healthineers. Jamadar is supported by an Australian National Health and Medical Research Council fellowship (APP1174164). Egan, Ward, and Jamadar are supported by the ARC Centre of Excellence for Integrative Brain Function (CE140100007).

## Data and Code Availability

The dataset containing the demographic, fMRI, PET, T1 structural and gradient field maps is freely available in BIDS format (Gorgolewski et al., 2016) from the *OpenNeuro* repository (http://openneuro.org) with the accession number ds002898 (Jamadar et al., 2020). Data and code that generated the results is available on the Open Science Network (DOI 10.17605/OSF.IO/DQN5S).

## Competing interests

The authors declare no competing interests.

## Notes

### Competing Interest Statement

The authors have declared no competing interest.

