## Appendix for "Metabolic and functional connectivity provide unique and complementary insights into cognition-connectome relationships"

### Supplementary Methods

#### Participants

**Table S1.** Participants' demographics (n = 26)

| | M (SD) | Range |
|---|---|---|
| Age (years) | 19.50 (1.36) | 18-23 |
| Gender (female/male) | 20(77%) / 6(23%) | |
| Education (years) | 14.77 (1.50) | 13-18 |
| Haemoglobin Levels (g/dl) | 15.25 (2.10) | 11.5-19.6 |

Note: M, Mean; SD, Standard deviation.

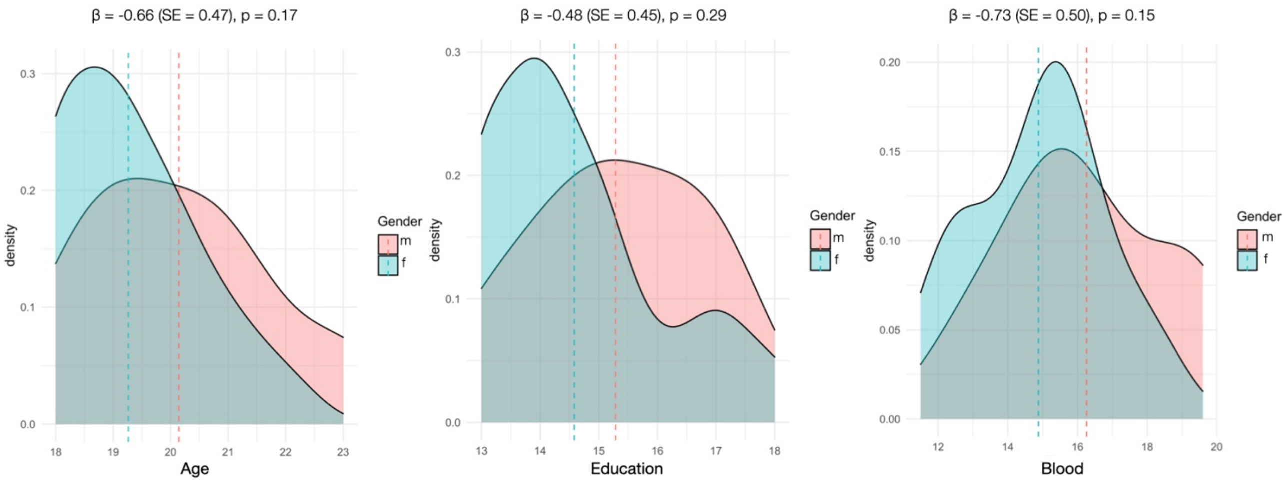


**Figure S1.** Gender as a function of age, education, and haemoglobin levels. Logistic regressions revealed that there were no gender effects across age (ß = -0.66, SE = 0.47, p = 0.17), education (ß = -0.48, SE = 0.45, p = 0.29) or haemoglobin levels (ß = -0.73, SE = 0.50, p = 0.15). For each logistic regression, the independent variable (i.e., age, education, and haemoglobin levels) was mean-centred, and the outcome variable (i.e., gender) was coded as male = 0, female = 1. ß = beta estimate, SE = standard error, p = significance level.

### Neuropsychological Test Battery

*Hopkins Verbal Learning Test-Revised*

The HVTL-R assess verbal episodic memory (Benedict et al., 1998). The HVTL-R consists of a 12-item word list drawn from three semantic categories (i.e., stones, animals and dwellings). Participants are asked to recall the list in any order after the list has been read out to them. This procedure is repeated three times. From this part of the test a total of the three recall trials (i.e., Total Recall score) is derived. The three learning trials are followed by a yes/no recognition trial, containing the 12 target words and 12 distracter words (6 semantically related and 6 semantically unrelated), from which a Recognition Score is derived. After a 20 – 25-minute delay, participants are asked to recall as many words as they can free-recall trial and a delayed recognition trial, from which a Delayed Recall and Retention Rate scores are derived, respectively.

*Symbol Digit Modalities Test*

The Symbol Digit Modalities Test (SDMT) is a neuropsychological test underpinned by divided attention, visual scanning, tracking and motor speed (i.e., processing speed, Smith, 1991). Participants are asked to pair specific numbers with given geometric figures using a reference key within 90 seconds. In total, there are nine possible pairs and participants have to complete 110 trials. The outcome variable is the number of correct, completed trials that were completed within 90 seconds (SDMT score).

*Stroop Neuropsychological Screening Test*

The Stroop Neuropsychological Screening Test (SNST) is a neuropsychological test assessing the executive function of inhibitory control, which occurs when the processing of a stimulus feature affects the simultaneous processing of another attribute of the same stimulus (Trenerry et al., 1989). Participants are asked to read two different tables as fast and accurately as possible. The first table (congruent Colour condition) consists of 112 colour names (i.e., red, green, blue, & tan) arranged in four columns of 28 names. The colour names are printed in different colour ink, but not in the matching colour (e.g., tan may be printed in red, but not tan). The second table (incongruent Colour-Word condition) is the same as the table for the Colour condition, except for the order of the colour names. For the congruent Colour condition, the individual is instructed to read the words aloud as quickly and accurately as she can. For the incongruent Colour-Word condition, the individual is instructed to name the colour of the ink in which the word is printed. Participant's completion time is measured via a stopwatch and 120 s are allowed for each stimulus sheet for a maximum test time of 4 min.

For the Colour-Word condition, participants are required to perform a less automated task (i.e., naming ink colour) while inhibiting the interference arising from a more automated task (i.e., reading the word; MacLeod & Dunbar, 1988). This difficulty in inhibiting the more automated process is called the Stroop effect (Stroop, 1935).While the SNST is widely used to measure the ability to inhibit cognitive interference; previous literature also reports its application to measure other cognitive functions such as attention, processing speed, cognitive flexibility (Jensen & Rohwer, 1966) and working memory (Engle and Kane, 2004). Thus, it may be possible to use the SNST to measure multiple cognitive functions.

We obtained four outcome variables from the SNST. First, the Incongruent Score, which is the number of correct responses completed minus incorrect responses during the incongruent Colour-Word condition. We inversed these values by multiplying them with -1 to ease interpretation: Larger negative values mean a better performance than smaller negative values. Second, the reaction time (RT) during the congruent Colour condition (i.e., congruent RT) time of completion for reading out the Colour table. Third, the RT during the incongruent Colour-Word condition (i.e., incongruent RT). And finally, a standardised score indicating frontal lobe dysfunction (Trenerry et al., 1989), whereby a value of 0 describes excellent functioning and a value of 1 describes frontal lobe dysfunction.

*Controlled Oral Word Association Test*

The Controlled Oral Word Association Test (COWAT) is a commonly used neuropsychological measure of phonemic fluency (Ruff et al., 1996). The COWAT has three-word conditions. The participant must produce as many words as she can that begin with a certain letter (F, A, or S) within one minute. The procedure is then repeated for the remaining two letters. Participants are instructed to exclude proper nouns, numbers, or the same word with a different suffix. Successful retrieval requires executive control over cognitive process such as selective attention, mental set shifting, internal response generation, and self-monitoring. The outcome variable is the total and correct number of word production.

*Colour Trails Task*

The Colour Trials (CT) Task (Dong et al., 2013) is a test to measure visuomotor tracking, scanning, divided attention and cognitive flexibility. The test is administered in two subcomponents: CT1 and CT2. During the first component, participants are presented with encircled numbers from 1 to 25 randomly distributed on a sheet of paper, and they are instructed to link the numbers in ascending order (i.e., 1-2-3…) using a pencil as fast and accurately as possible. During the second component, participants are asked still asked to link the numbers from 1 to 25 in ascending order as fast and accurately as possible but alternating the order respective of the numbers' (pink and yellow). Numbers are shown in pink and yellow for both task components, but participants are inly made aware of the colouring during the second component. For example, the participants must trace the line from pink number 1 to yellow number 2, to pink number 3 and so on. Task performance for each part is quantified by measuring the correct scores for each part (CT1 and CT2), as well as an

Interference Index, which indicates the difference in time needed to complete CT2 as opposed to CT1.

*Center for Epidemiologic Studies Depression Scale – Revised*

As part of our cognitive battery, we also assessed participants depression scores via the Center for Epidemiologic Studies Depression Scale – Revised (CESD-R; Radloff, 1977). This scale is a 20 item self-report questionnaire used to measure symptoms of depression over time. The instrument measures nine different symptom groups of Major Depression Disorder as defined by the American Psychiatric Association Diagnostic and Statistical Manual (DSM-V): dysphoria, anhedonia, appetite, sleep, concentration, guilt, fatigue, agitation, suicidal ideation. The outcome variable is the CESD-R score, which is calculated by finding the sum of 20 items and ranges from 0-60. A score equal to or above 16 indicates a person at risk for clinical depression.

### Supplementary Results

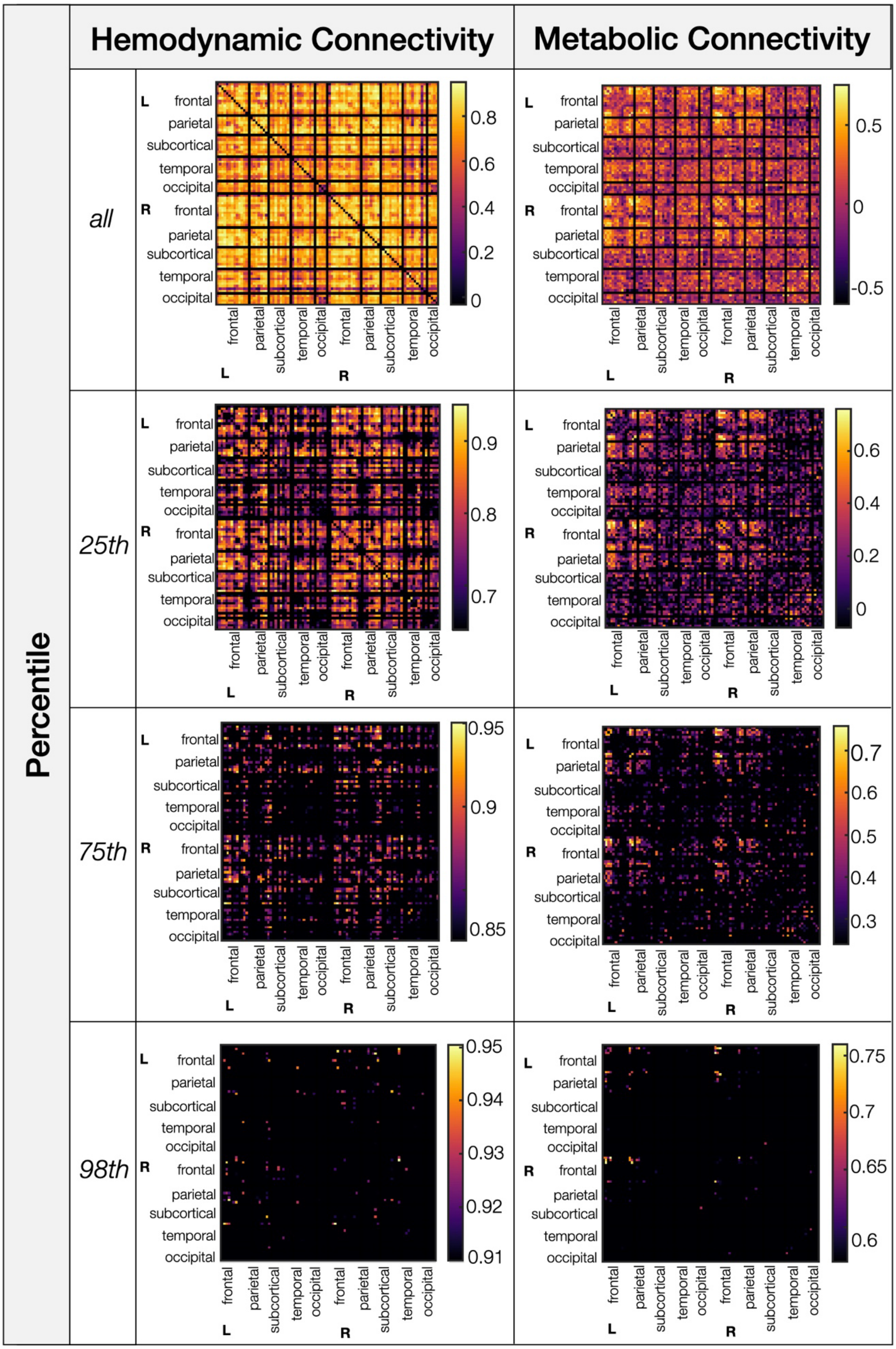

**Figure S2.** Hemodynamic and Metabolic Connectivity loadings. Loadings were thresholded, so that all loadings (first row), loadings of the 25th percentile (second row), loadings of the 75th percentile (third row), and loadings for the 98th percentile (last row) are shown. L, Left; R, Right.

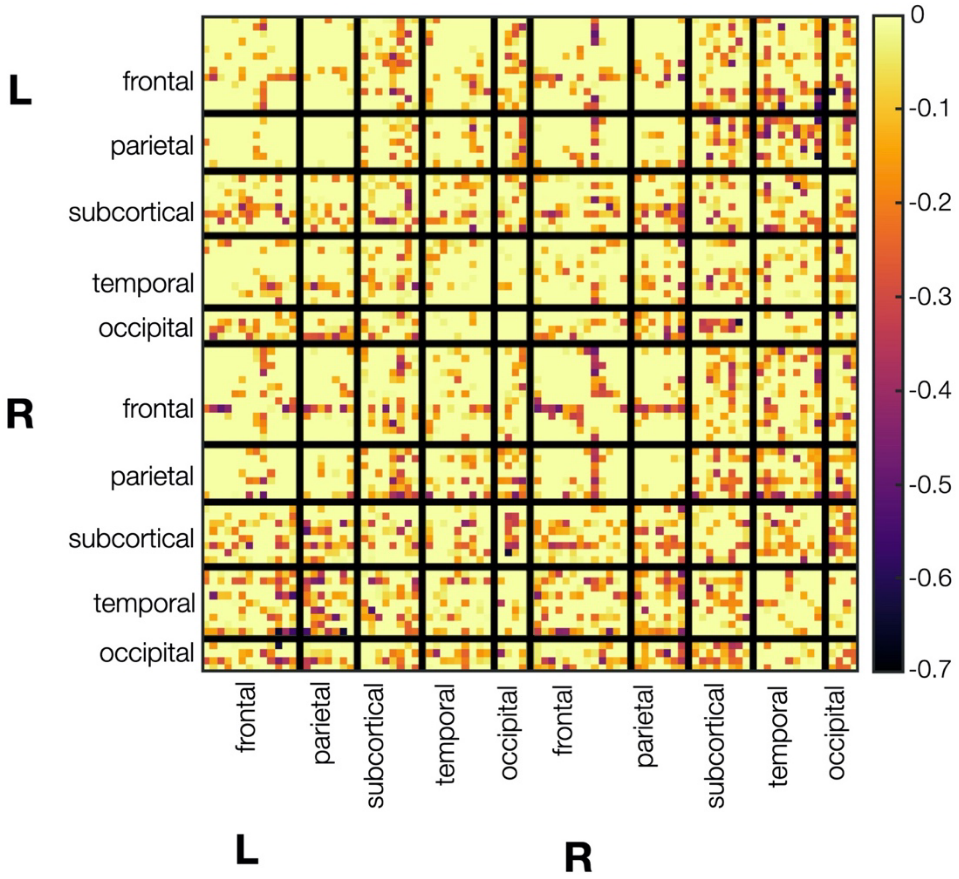


**Figure S2.** Negative metabolic connectivity loadings. L, left; R, Right.
